# Differential Expression of Gluconeogenesis-Related Transcripts in a Freshwater Zooplankton Model Organism Suggests a Role of the Cori Cycle in Hypoxia Tolerance

**DOI:** 10.1101/2023.04.06.535910

**Authors:** M. C. Malek, J.R. Behera, A. Kilaru, L. Y. Yampolsky

## Abstract

1. Gluconeogenesis (GNG) is the process of regenerating glucose and NAD+ that allows continuing ATP synthesis by glycolysis during fasting or in hypoxia. Recent data from *C. elegans* and crustaceans challenged with hypoxia show differential and tissue-specific expression of GNG-specific genes.
2. Here we report differential expression of several GNG-specific genes in the head and body of a model organism, *Daphnia magna,* a planktonic crustacean, in normoxic and acute hypoxic conditions. We predict that GNG-specific transcripts will be enriched in the body, where most of the fat tissue is located, rather than in the head, where the tissues critical for survival in hypoxia, the central nervous system and locomotory muscles, are located. We measured the relative expression of GNG-specific transcripts in each body part by qRT-PCR and normalized them by either the expression of a reference gene or the rate-limiting glycolysis enzyme pyruvate kinase (PK).
3. Our data show that of the three GNG-specific transcripts tested, pyruvate carboxylase (PC) showed no differential expression in either the head or body. Phosphoenolpyruvate carboxykinase (PEPCK-C), on the other hand, is upregulated in hypoxia in both body parts. Fructose-1,6-bisphosphatase (FBP) is upregulated in the body relative to the head and upregulated in hypoxia relative to normoxia, with a stronger body effect in hypoxia when normalized by PK expression.
4. These results support our hypothesis that *Daphnia* can survive hypoxic conditions by implementing the Cori cycle, where body tissues supply glucose and NAD+ to the brain and muscles, enabling them to continuously generate ATP by glycolysis.

## Introduction

The role of the Cori cycle or gluconeogenesis (GNG) has been well characterized in humans and other mammals, where products of glycolysis are utilized in the liver to resupply muscles and other critical tissues with glucose and NAD+, thus allowing glycolysis to continuously generate ATP during bursts of muscular activity or while in hypoxic or fasting conditions [1–6]. In some other vertebrates adapted to either periodic fasting[7] or hypoxia[8], the Cori cycle emerges as a key element of adaptation to environmental extremes. The GNG pathway utilizes lactate generated through the anaerobic catabolism of pyruvate as a precursor for glucose synthesis. This multi-step process is assisted by the respective glycolytic enzymes catalyzing the reverse reactions, except for the exclusive involvement of phosphoenolpyruvate carboxykinase (cytosolic PEPCK-C and mitochondrial PEPCK-M), fructose 1,6-bisphosphatase (FBP), and glucose 6-phosphatase in GNG[6] (Fig. 1). The upregulation of GNG-specific genes, primarily PEPCK paralogs, in hypoxia and the role of hypoxia-inducible factor (HIF-1) in this upregulation have been reliably demonstrated in mammalian models[9–11].

**Fig. 1.**
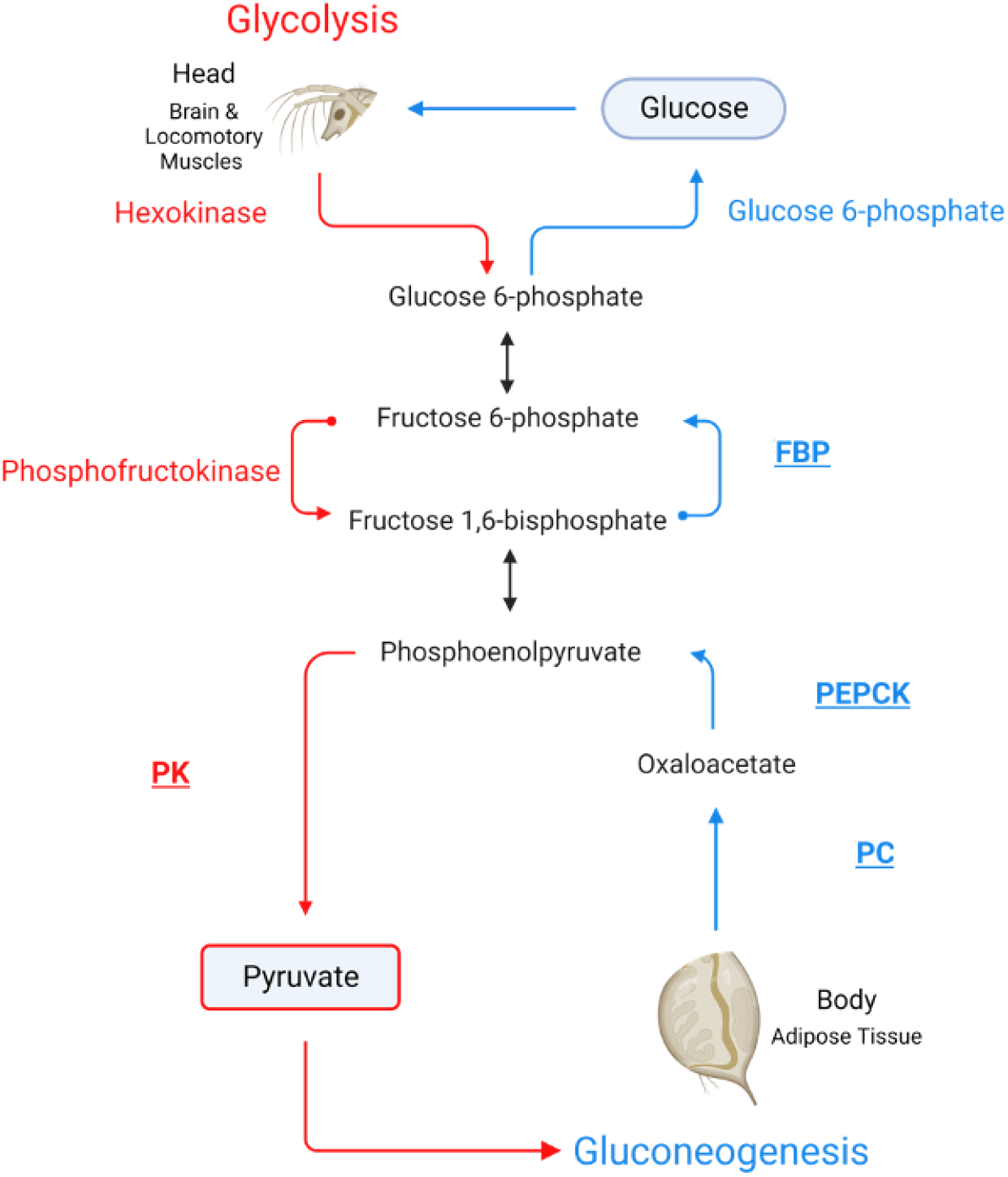
Schematic representation of the Cori cycle. Only those steps of glycolysis (red) and gluconeogenesis (blue) that are accomplished by different enzymes are labeled. The enzymes labeled are rate-limiting for each of the two branches of the cycle. Enzymes analyzed in this study are shown in bold, underlined type. FBP: fructose-1,6-bisphosphatase; PC: pyruvate carboxylase; PEPCK: phosphoenolpyruvate carboxykinase; PK: pyruvate kinase.

However, little data is available on GNG and the Cori cycle in aquatic invertebrates like crustaceans. Aquatic organisms are more likely to experience periods of hypoxia than terrestrial ones, as oxygen solubility and diffusion rates in water are low, and its availability is highly dependent on temperature, the respiratory activity of aerobic heterotrophs, and the non-biological oxidation of organic matter. Therefore, it is important to examine GNG as a possible mechanism of hypoxic survival to understand the ability of organisms to cope with episodes of high temperatures and low oxygen availability in their natural habitats.

Studies on the decapod shrimp *Litopenaeus vannamei* indicate tissue-specific expression of GNG-related genes are consistent with the role of GNG in providing glucose to fuel muscles during hypoxia. The hepatopancreas in crustaceans, an organ that is functionally analogous to the vertebrate liver, shows higher expression of *pyruvate carboxylase (PC)* [12], PEPCK-C and PEPCK-M [13], and FBP [14,15] relative to muscle or gill tissues. Importantly, this tissue-specific gene expression is further enhanced by hypoxia [12,13,15]. Furthermore, glucose-6-phosphatase, one of the rate-limiting enzymes of glycolysis, shows a reverse response pattern to hypoxia, where it is upregulated in the gills but not in the hepatopancreas [16]. These results are consistent with the operation of the Cori cycle to allow hypoxia tolerance, with the hepatopancreas completing the GNG phase of the cycle and supplying tissues like muscles with glucose and NAD+. We expect this mechanism is likely conserved across invertebrates, as a similar differential expression was observed in the nematode *Caenorhabditis elegans* [17].

We have previously analyzed differential gene expression in response to mild chronic and severe acute hypoxia in a classic model organism for aquatic ecophysiology, *Daphnia magna* [18] revealing that while only a small subset of abundant transcripts showed differential expression responses under mild chronic hypoxic conditions, many, including the GNG-specific PEPCK-C, showed upregulation in acute severe hypoxia. Aralar1, a mitochondrial carrier protein that transports aspartate from the mitochondria to the cytosol, also showed upregulation in hypoxia. The function of Aralar1 is critical for GNG, as it allows bypassing pyruvate as the starting point of the pathway by converting aspartate to oxaloacetate, a PEPCK-C substrate that converts to phosphoenolpyruvate in the cytosol.

The goal of this study was to examine the differential expression of GNG-related enzymes PC, PEPCK-C, and FBP and the Aralar1 transporter separately in the head and body of hypoxia-challenged vs. normoxic control *Daphnia*. For *Daphnia* to survive in an acute hypoxic state, metabolic processes must be altered to allow alternative sources of glucose through GNG. We hypothesize that if GNG is upregulated as part of the Cori cycle, the relative abundance of transcripts should correspond to either the head or the body, with the body containing the majority of GNG transcripts while the head undergoes glycolysis steps (Figure 1). To that end, we used both a reference gene unrelated to glucose metabolism and a rate-limiting glycolysis enzyme, pyruvate kinase, as normalization controls to infer the relative expression of GNG-related transcripts, thus emphasizing the relative expression of the two opposing Cori cycle pathways in the heads and bodies of *Daphnia*.

## Materials and methods

### Daphnia clones and culture

*Daphnia magna* stocks used in this study were obtained from the Basel University Daphnia Stock Collection (Basel, Switzerland) and maintained locally since 2016. The IDs of the four stocks used were FI-FSP1-16-2, GB-EL75-69, HU-K-6, and IL-M1-8; details of the provenance of these *D. magna* clones are supplied on Table 1 in Ekwudo et al. 2022. Hereafter, we refer to these clones by the first two letters of their Basel stock IDs. Previous longevity studies indicate that these clones differ significantly in their lifespan and acute hypoxia tolerance, with FI and IL being the short-lived, hypoxia-tolerant clones, and GB and HU showing higher longevity but lower hypoxia tolerance [18]. Stocks were maintained in modified ADaM zooplankton medium (Ref. [19]; https://evolution.unibas.ch/ebert/lab/adam.htm) at the density of one adult Daphnia per 20 mL, at 20 °C under 16h:8h L:D photoperiod and fed green alga *Scenedesmus acutus* Meyen (current nomenclature *Tetradesmus obliquus* (Turpin) M. J. Wynne) at the concentration of 10^5^ cells per mL per day. Newborn individuals of each of the four clones were collected within 24 h of birth and placed in groups of 10 in 100 mL jars, with the density reduced to one Daphnia per 20 mL at maturity (day 6-8), and maintained with food added daily, water changed, and neonates removed twice weekly. The four clones used were characterized by their lactate:pyruvate ratio in normoxic and acute hypoxic conditions, as described below.

**Table 1:**
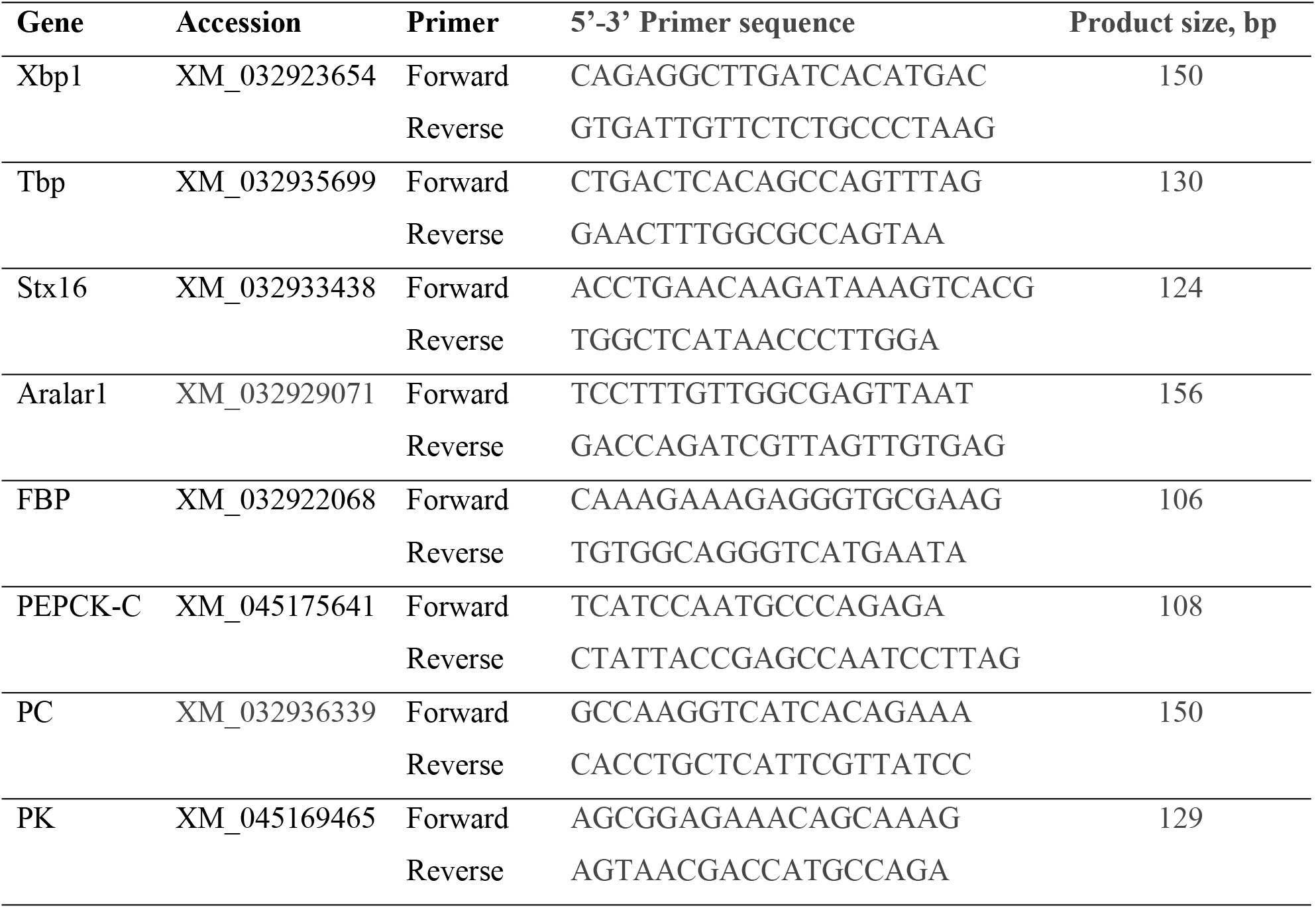
Gene-specific primers used in qRT-PCR.

### Acute hypoxia exposure and lactate:pyruvate ratio measurements

*Daphnia* females maintained as described above until the age of 15-25 days were randomly assigned to control and hypoxia treatments. Hypoxia treatment individuals were placed individually in 10-mL screw-cap vials filled with 20 °C ADaM water deoxygenized to <1 mg O_2_/L by intense bubbling with nitrogen. Oxygen concentration was maintained by an Extech DO210 dissolved oxygen meter. Simultaneously, the control *Daphnia* were transferred into fresh vials containing fully oxygenized (>8 mg O_2_/L) ADaM. No food was added to either the hypoxia or control vials. *Daphnia* were harvested after 12 hours of exposure. Typically, no mortality occurs in 12 hours at 1 mg O_2_/L, with survival times ranging between 17 and 30 hours[18]. This experiment was conducted in two randomized blocks on two different dates, with 48 and 16 *Daphnia* used in each block.

Whole *Daphnia* females from either control or acute hypoxia treatments were homogenized in 100 μL of RO water using a bead beater with 0.15 mm ZrO beads and centrifuged for 4 min at 4 °C. The supernatant was then used to determine lactate and pyruvate concentrations using CellBioLabs® colorimetric kits (Cat. #s MET-5012 and MET-5125, respectively), according to the manufacturer’s protocol, scaled down to 50 μL reactions, each containing 20 μL of supernatant. Absorbance was measured in 384-well plates on a BioTek Synergy plate reader at room temperature at 490 and 570 nm for lactate and pyruvate, respectively. In parallel, soluble protein content in the supernatant was measured by the Bradford method to normalize lactate and pyruvate concentrations per mg of protein.

Based on the results, we then selected the hypoxia-tolerant clone IL, characterized by the highest lactate:pyruvate ratio in hypoxia, for further qRT-PCR analyses.

### Differential gene expression quantification

Total RNA was extracted using the RNeasy Mini Kit (Qiagen) from the heads and bodies of 15 days old *D. magna* females (clone IL) exposed to acute hypoxia treatment or normoxia as described above. 10 females per sample were used for the RNA extraction, and four such samples were prepared for each condition. The RNA concentration was measured by Nanodrop, and samples were diluted to a final concentration of 20 ng/μL in RNase-free water. qRT-PCR was done using the qScript One-Step SYBR Green qRT-PCR Kit (Quantabio) according to the manufacturer’s protocol, using gene-specific primers (Table 1). A total of 80 ng of RNA was used as input in a 20 μL reaction mixture, and all the reactions were carried out using an Illumina Eco Real-time PCR system. Four biological replicate samples in two technical replicates were used. The cDNA synthesis was performed at 50 °C for 10 minutes using the primers specific for the genes (Table 1), followed by polymerase activation at 95 ^°^C for 10 minutes. PCR cycles were carried out at 95 ^°^C for 10s, followed by 60 °C for 20s, for 40 cycles. Fluorescence signals (Channel 1; for SYBR Green) were detected after each cycle. The reference genes were chosen following Ref[20]. We tested *D. magna* orthologs of the following proposed internal reference genes for constant expression: X-box binding protein 1 (*Xbp1*), TATA-box binding protein (*Tbp*), and syntaxin-16 (*Stx16*). Among these,*Xbp1* showed no difference in expression between the control and hypoxia treatments and was thus chosen as the reference gene. The quantification cycle (Cq) values for each gene expression were determined from the amplification curves at a threshold value set at 0.02, and the relative expression of GNG-related genes was expressed as - ΔCq (Rao et al. 2013). For further comparison relative to the pace of glycolysis, corresponding gene expressions were also normalized against that of the muscular isoform pyruvate kinase (PK) gene that catalyzes one of the rate-limiting steps of glycolysis.

Table 1. qRT-PCR primers used for GNG-related and reference genes. Xbp1, X-box binding protein 1; Tbp, TATA-box binding protein; Stx16, syntaxin-16; Arlar1; FBP, fructose 1,6-bisphosphotase; PEPCK-C, cytoplasmic phosphoenolpyruvate carboxykinase; PC, mitochondrial pyruvate carboxylase; PK, pyruvate kinase, muscle isoform.

### Data analysis

Lactate and pyruvate content and qRT-PCR data were analyzed using Residual Maximum Likelihood (REML) ANOVA using JMP (Ver. 16, SAS Institute 2016), with protein-normalized lactate and pyruvate concentrations and their ratio, or relative expression, respectively, as the response variables. The main effects in the model were clones and acute hypoxia exposure. The date of measurement was included in the model as a random block effect. For qRT-PCR data, the biological replicate was included in the analysis as a random block effect. Table-wide sequential Bonferroni-adjusted P-values[20] were calculated for each table of results.

## Results

### Lactate and pyruvate content

Here we examined the differences in protein-normalized concentrations of lactate and pyruvate and their ratio under hypoxic and normoxic conditions among four *D. magna* clones (Table 2, Fig 2). Lactate concentration significantly increased in hypoxia but did not differ among the clones, while pyruvate concentration did not increase in hypoxia but showed a strong clone effect, with the IL clone demonstrating the lowest levels of pyruvate. As the results indicate, the lactate:pyruvate ratio showed both clone and hypoxia effects, with the increased lactate:pyruvate ratio after exposure to acute hypoxia being largely due to the increase in lactate concentration, whereas interclonal differences were largely due to differences in pyruvate concentration. There were no interaction effects between clones and hypoxia in either lactate or pyruvate concentrations or their ratio (Table 2).

**Table 2.** Residual maximum likelihood (REML) analysis of variance of the differences in protein-normalized concentrations of lactate and pyruvate and their ratio among clones and between hypoxia treatment and normoxic control. P<0.025 shown in italics; P<0.001 in bold. Sequential Bonferroni-corrected P-values are should to the right of individually significant P-values

| Source | DF | DFDen | F Ratio | P | P<sub>adj</sub> |
| --- | --- | --- | --- | --- | --- |
| **Response Lac/Pyr** |  |  |  |  |  |
| Clone | 3 | 55 | 8.69 | **<0.0001** | **<0.001** |
| Hypoxia | 1 | 47.02 | 25.43 | **<0.0001** | **<0.001** |
| Clone*Hypoxia | 3 | 55 | 1.27 | 0.29 |  |
| **Response Lac, mM/mg Prot** |  |  |  |  |  |
| Clone | 3 | 55 | 0.46 | 0.71 |  |
| Hypoxia | 1 | 55.9 | 63.47 | **<0.0001** | **<0.001** |
| Clone*Hypoxia | 3 | 55 | 0.397 | 0.75 |  |
| **Response Pyr, mM/mg Prot** |  |  |  |  |  |
| Clone | 3 | 55 | 7.02 | **0.0004** | **0.0024** |
| Hypoxia | 1 | 55.89 | 6.18 | *0.0159* | 0.08 |
| Clone*Hypoxia | 3 | 55 | 0.08 | 0.97 |  |

**Fig. 2.**
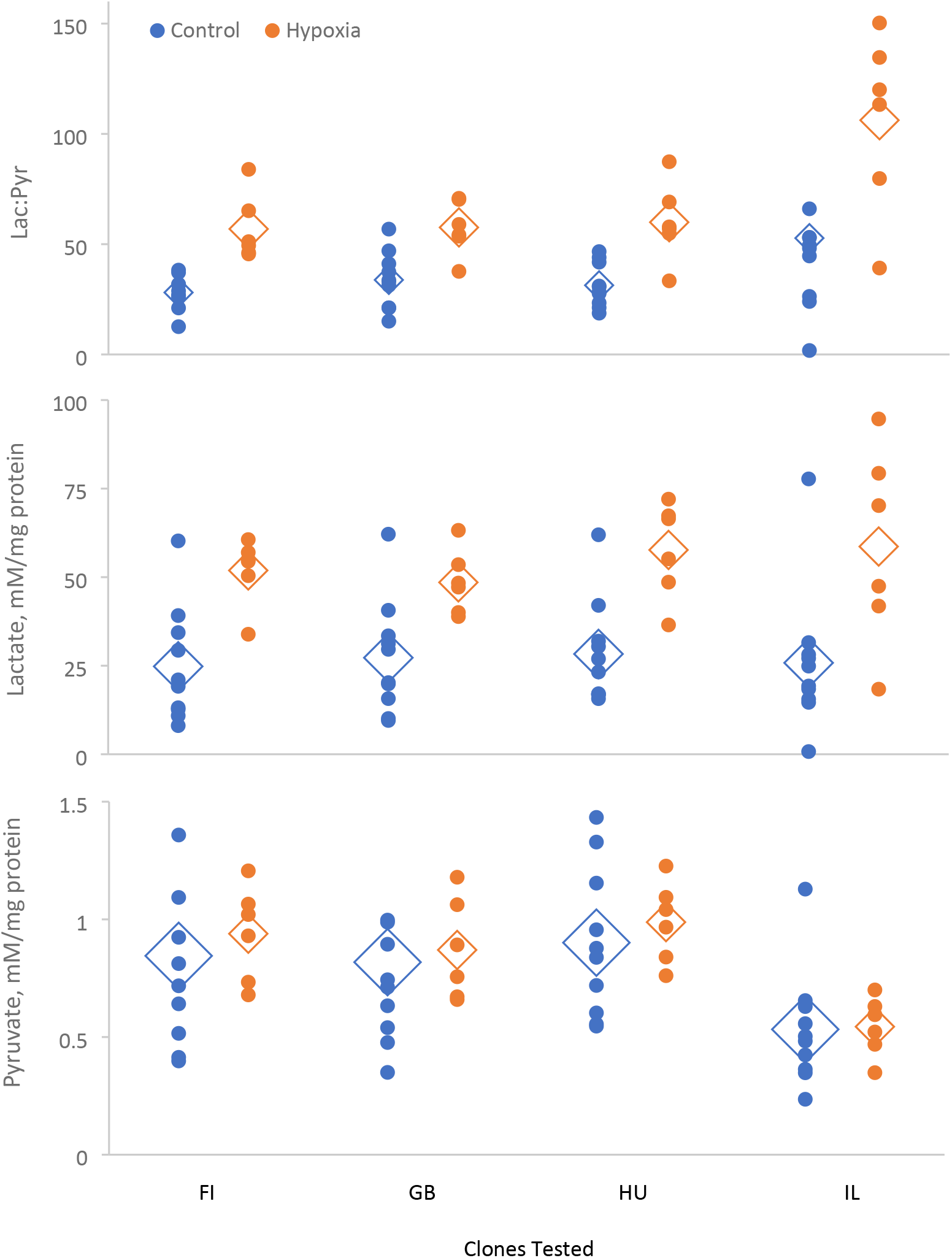
Lactate:pyruvate ratio and protein-normalized lactate and pyruvate concentration in whole-body extracts in Daphnia from normoxic control (blue) and 12 h exposure to acute hypoxia (orange). Diamonds represent means; a diamond’s height represents the SE of the mean.

### Differential gene expression

The results of differential expression analysis by qPCR were somewhat different depending on which transcript was used for normalization: a carbohydrate metabolism-independent housekeeping reference gene Xbp1 or the glycolysis rate-limiting PK (Fig. 1), even though PK itself showed no significant differences between either oxygen levels or body parts (Table 3, Fig. 3). Normalization of GNG-specific transcripts by Xbp1 reflects general levels of expression; normalization by PK reflects the relative activity of the GNG vs. glycolysis branches of the Cori cycle. Of the four GNG-related transcripts tested, the transport protein Aralar1 transcript, contrary to predictions, showed a slight downregulation in hypoxia. PC showed no evidence of differential expression regardless of which reference was used for normalization (Table 3). The other two transcripts, PEPCK-C and FBP, showed hypoxia-related differential expression. Both PEPCK-C and FBP were upregulated in hypoxia when normalized by Xbp1 (tentatively significant after multiple test correction); FBP was both upregulated in hypoxia and in the bodies, relative to the heads, as predicted, with the hypoxia upregulation being stronger in the head than in the body (Table 3; Fig. 3).

**Table 3.**
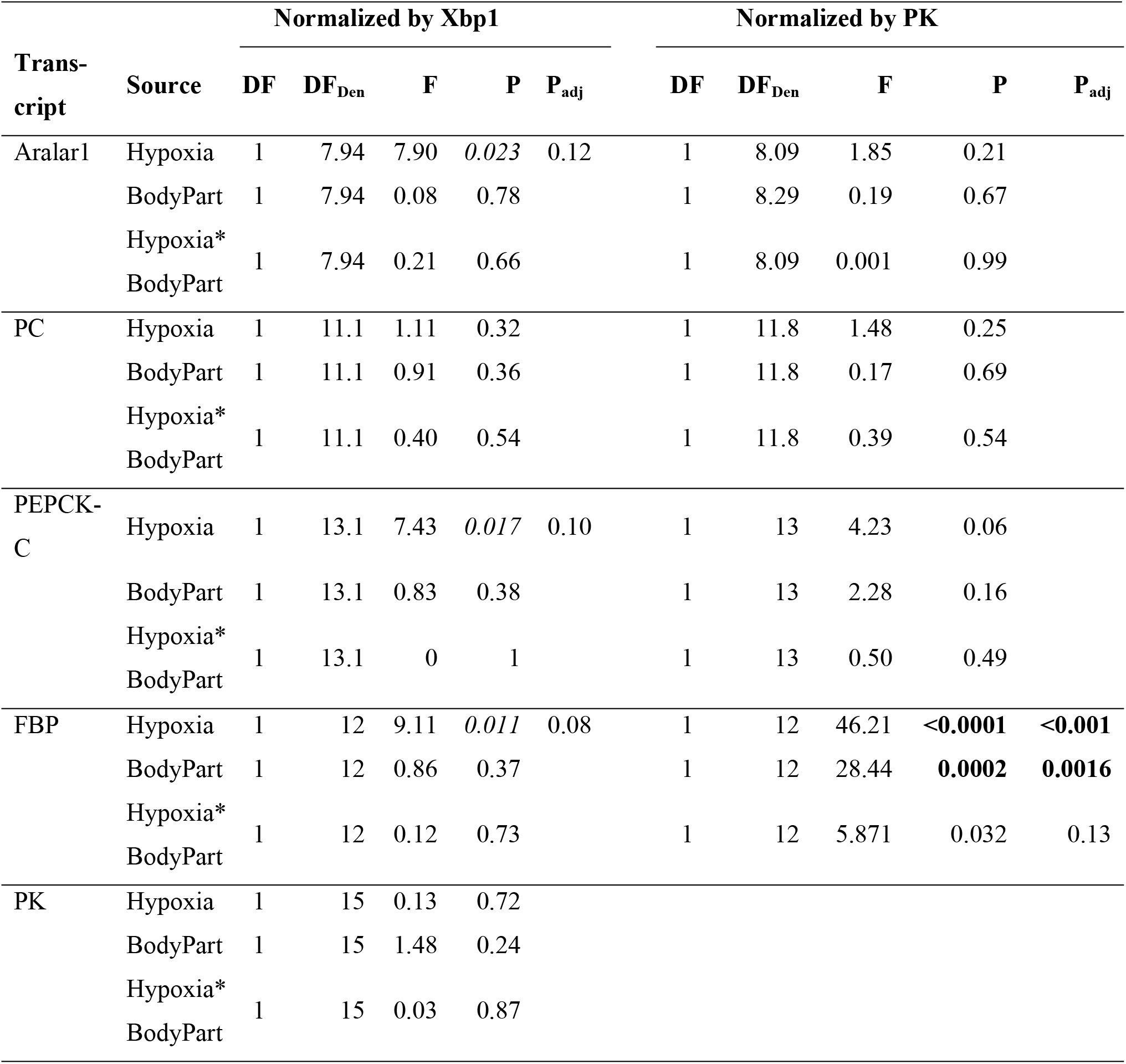
Residual maximum likelihood (REML) analysis of variance of the effects of hypoxia (normoxic control vs. 12 hours at <1 mg/L O_2_) and body part (head vs. body) on transcript abundance (-ΔCq) for four GNG-related transcripts normalized either by the general housekeeping reference gene encoding X-box binding protein (Xbp1) or glycolysis-specific pyruvate kinase (PK). P<0.025 shown in italics; P<0.001 in bold. Sequential Bonferroni-adjusted P-values shown on the right on individually significant P-values.

**Fig. 3.**
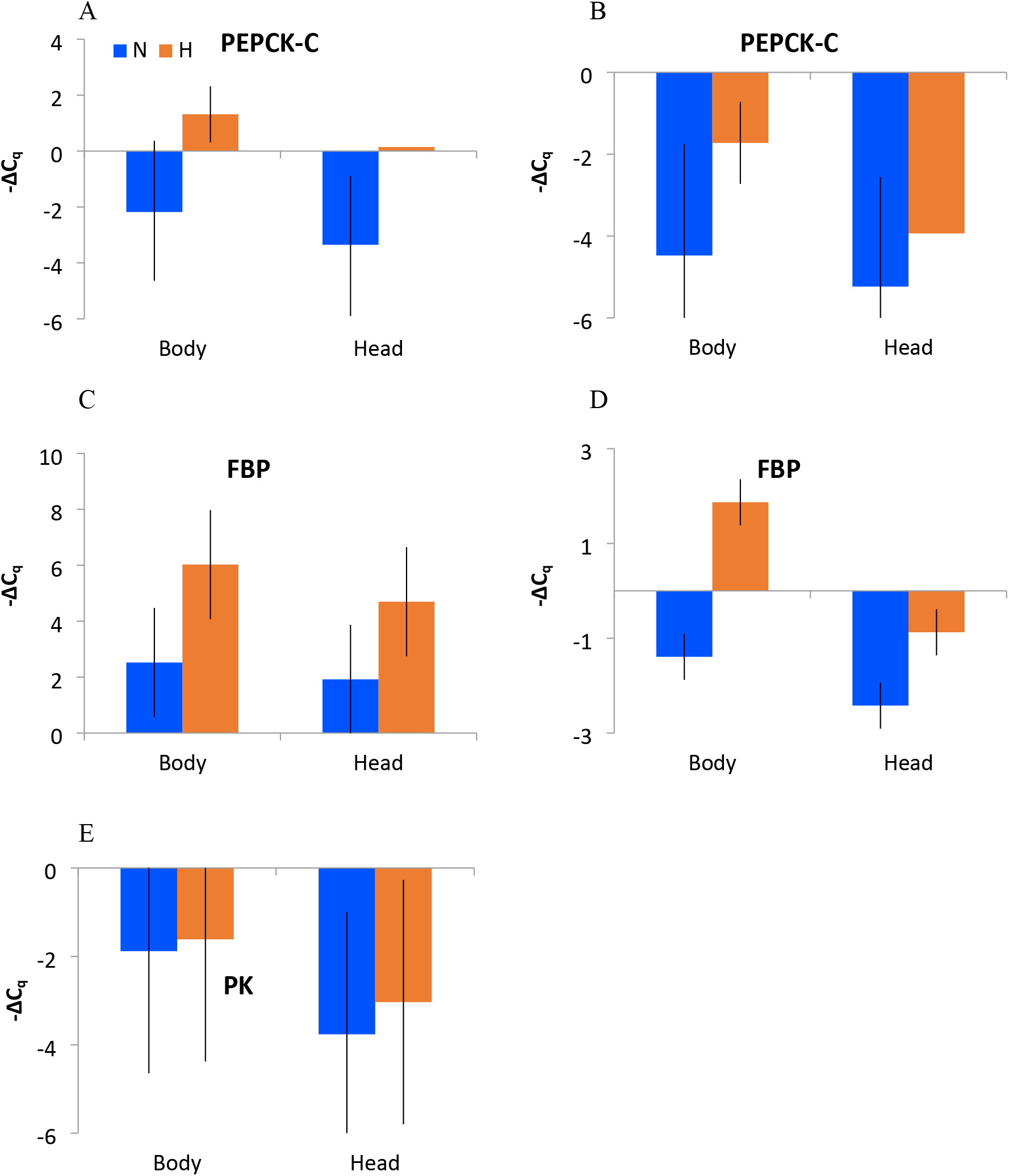
Relative expression (-ΔC_q_) of PEPCK-C (A, B), FBP (C, D), and PK (E) transcripts normalized by Xbp (A, C, E) and PK (B, D). Error bars represent standard error. Higher values indicate higher expression; -ΔC_q_ difference of 1 corresponds to two-fold difference in transcript abundance. Colors as on Fig. 2.

## Discussion

The Cori cycle is an adaptation to intense bouts of muscular activity in larger animals, where it is difficult to supply sufficient oxygen for oxidative phosphorylation in muscles during peak ATP demand [5]. However, this is a costly way of generating ATP, as for every mole of ATP generated by glycolysis in the muscles, three moles are spent in the GNG branch of Cori cycle, making it unsustainable in the long term. However, even small organisms may have to resort to GNG to regenerate glucose in hypoxic conditions. For example, zooplankton organisms like *Daphnia* must continuously swim (typically upward) to avoid hypoxic layers or spots within a lake or pond, and this is likely a subject for selection.

In *Daphnia* and other cladocerans, the head contains major locomotory muscles, namely the ones driving the 2^nd^ (swimming) antenna, in addition to the mitochondria-rich ATP-consuming central neural system (Ref. [21], Fig. 1.3). Thus, the head tissues and organs are likely to be the primary consumers of glucose synthesized by GNG in conditions when glycolysis is the main source of ATP. While the “classic” mammalian GNG-active organs, the liver and kidneys, do not have direct counterparts in *Daphnia* anatomy, we hypothesize that the adipose tissue, located in the thorax and the abdomen, may be the main site of GNG. Lactate and pyruvate metabolism, including the starting points of gluconeogenesis such as the conversion of pyruvate to oxaloacetate by PC, is known to be active in adipose tissue in both mammals and arthropods [22]. Therefore, we hypothesize that the “body” (thorax and abdomen) may be the donor of glucose produced by GNG in the Cori cycle operating in *Daphnia* under hypoxic conditions, suggesting differential expression of the GNG and glycolysis-related genes in the body and the head. And a further prediction made was that acute hypoxia should augment this differentiation.

These predictions were tested in a qRT-PCR experiment, measuring GNG-related transcript abundance separately in the heads and bodies of individuals from the IL *Daphnia* clone. We selected this clone from a panel of four as it was the one characterized by the lowest pyruvate accumulation in tissues, hypothetically indicating the utilization of pyruvate for lactic fermentation and/or GNG. This clone has previously been demonstrated to be the most hypoxia-tolerant of the four clones tested [18]. We observed a significant upregulation of PEPCK-C and FBP (but not of PC) in hypoxia in both body parts when normalized by a reference gene unrelated to carbohydrate metabolism (Fig. 3). We also observed a significant upregulation of FBP, a rate-limiting GNG enzyme, in hypoxia and in the body relative to the head, when normalized by PK, one of the rate-limiting enzymes of glycolysis. We therefore conclude that body/head compartmentalization of the glycolysis and GNG pathways is likely necessary for survival in hypoxia. Furthermore, we predicted the hypoxia-by-body part interaction with the hypoxia effect being stronger in the body than in the head (Fig. 3); however, this predicted interaction effect was only marginally significant and did not survive multiple test correction.

It is yet to be tested if the same observation would be made with a less hypoxia-tolerant clone of *D. magna* (or in less hypoxia-tolerant species of zooplankton). Further studies would test if that variation in hypoxia tolerance, within and among zooplankton species, is maintained by the trade-offs between the ability to operate the Cori cycle in hypoxia and the GNG-associated costs of doing so when oxygen is abundant.

## Conclusions

We observe upregulation of the genes encoding rate-limiting gluconeogenesis enzymes, cytoplasmic phosphoenolpyruvate carboxykinase (PEPCK-C) and fructose-1,6-bisphosphatase (FBP), in hypoxia within a hypoxia-tolerant clone of *Daphnia.* When normalized by the rate-limiting glycolysis enzyme pyruvate kinase, the upregulation of FBP is more pronounced in the body than in the head of *Daphnia,* indicating the potential role of the Cori cycle in sustaining glycolysis in the central nervous system and locomotory muscles during hypoxia.

## References

1. Cori, C.F., & Cori, G.T. 1929. Glycogen formation in the liver from d- and l-lactic acid. J Biol Chem, 81, 389–403.

2. Reichard, G. A., Jr., Moury N. F., Jr., Hochella, N. J., Patterson, A. L., & Weinhouse, S. 1963. Quantitative estimation of the Cori cycle in the human. J. Biol. Chem, 238, 495.

3. Cahill, G. F. 1970. Starvation in Man. New England Journal of Medicine, 282(12), 668–675. doi: 10.1056/nejm197003192821209

4. Freminet, A., Poyart, C., Leclerc, L., & Gentil, M. 1976. Effect of fasting on the Cori cycle in rats. FEBS Lett, 66, 2, 328–331. doi: 10.1016/0014-5793(76)80532-7. PMID: 955096.

5. Cori, C.F. 1981. The glucose-lactic acid cycle and gluconeogenesis. Current Topics in Cellular Regulation, 18, 377–387.

6. Nelson, D.L., & Cox, M.M. 2005. Lehninger Principles of Biochemistry (Fourth ed.). New York: W.H. Freeman and Company. p. 543.

7. Champagne, C.D., Houser, D.S., & Crocker, D.E. 2005. Glucose production and substrate cycle activity in a fasting adapted animal, the northern elephant seal. J Exp Biol., 208(Pt 5), 859–868. doi: 10.1242/jeb.01476. PMID: 15755884.

8. Gupta, A., Varma, A., & Storey, K.B. 2021. New insights to regulation of fructose-1,6-bisphosphatase during anoxia in red-eared slider, Trachemys scripta elegans. Biomolecules, 11, 10, 1548. doi: 10.3390/biom11101548. PMID: 34680181.

9. Choi, J.H., Park, M.J., Kim, K.W., Choi, Y.H., Park, S.H., An, W.G., Yang, U.S., & Cheong J. 2005. Molecular mechanism of hypoxia-mediated hepatic gluconeogenesis by transcriptional regulation. FEBS Lett., 579(13), 2795–2801. doi: 10.1016/j.febslet.2005.03.097. PMID: 15907483.

10. Hara, Y., & Watanabe, N. 2020. Changes in expression of genes related to glucose metabolism in liver and skeletal muscle of rats exposed to acute hypoxia. Heliyon, 6, 7, e04334. doi: 10.1016/j.heliyon.2020.e04334. PMID: 32642586.

11. Owczarek, A., Gieczewska, K., Jarzyna, R., Jagielski, A.K., Kiersztan, A., Gruza, A., & Winiarska, K. 2020. Hypoxia increases the rate of renal gluconeogenesis via hypoxia-inducible factor-1-dependent activation of phosphoenolpyruvate carboxykinase expression. Biochimie, 171-172, 31–37. doi: 10.1016/j.biochi.2020.02.002. PMID: 32045650.

12. Granillo-Luna, O.N., Hernandez-Aguirre, L.E., Peregrino-Uriarte, A.B., Duarte-Gutierrez, J., Contreras-Vergara, C.A., Gollas-Galvan, T., & Yepiz-Plascencia, G. 2022. The anaplerotic pyruvate carboxylase from white shrimp Litopenaeus vannamei: Gene structure, molecular characterization, protein modelling and expression during hypoxia. Comp Biochem Physiol A Mol Integr Physiol, 269, 111212. doi: 10.1016/j.cbpa.2022.111212. PMID: 35417748.

13. Reyes-Ramos CA, Peregrino-Uriarte AB, Cota-Ruiz K, Valenzuela-Soto EM, Leyva-Carrillo L, Yepiz-Plascencia G. 2018. Phosphoenolpyruvate carboxykinase cytosolic and mitochondrial isoforms are expressed and active during hypoxia in the white shrimp Litopenaeus vannamei. Comp Biochem Physiol B Biochem Mol Biol. 226:1–9. doi: 10.1016/j.cbpb.2018.08.001. PMID: 30107223.

14. Cota-Ruiz, K., Peregrino-Uriarte, A.B., Felix-Portillo, M., Martínez-Quintana, J.A., & Yepiz-Plascencia, G. 2015. Expression of fructose 1,6-bisphosphatase and phosphofructokinase is induced in hepatopancreas of the white shrimp Litopenaeus vannamei by hypoxia. Mar Environ Res, 106, 1–9. doi: 10.1016/j.marenvres.2015.02.003. PMID: 25725474.

15. Cota-Ruiz, K., Leyva-Carrillo, L., Peregrino-Uriarte, A.B., Valenzuela-Soto, E.M., Gollas-Galván, T., Gómez-Jiménez, S., Hernández, J., & Yepiz-Plascencia, G. 2016. Role of HIF-1 on phosphofructokinase and fructose 1,6-bisphosphatase expression during hypoxia in the white shrimp i. Comp Biochem Physiol A Mol Integr Physiol, 198, 1–7. doi: 10.1016/j.cbpa.2016.03.015. PMID: 27032338.

16. Hernández-Aguirre, L.E., Cota-Ruiz, K., Peregrino-Uriarte, A.B., Gómez-Jiménez, S., & Yepiz-Plascencia, G. 2021. The gluconeogenic glucose-6-phosphatase gene is expressed during oxygen-limited conditions in the white shrimp Penaeus (Litopenaeus) vannamei: Molecular cloning, membrane protein modeling and transcript modulation in gills and hepatopancreas. J Bioenerg Biomembr, 53, 4, 449–461. doi: 10.1007/s10863-021-09903-6. PMID: 34043143.

17. Vora, M., Pyonteck, S.M., Popovitchenko, T., Matlack, T.L., Prashar, A., Kane, N.S., Favate, J., Shah, P., & Rongo, C. 2022. The hypoxia response pathway promotes PEP carboxykinase and gluconeogenesis in C. elegans. Nature Communications, 13, 1, 6168. doi: 10.1038/s41467-022-33849-x. PMID: 36257965.

18. Ekwudo MN, Malek MC, Anderson CE, Yampolsky LY. 2022. The interplay between prior selection, mild intermittent exposure, and acute severe exposure in phenotypic and transcriptional response to hypoxia. Ecol Evol. 12(10):e9319. doi: 10.1002/ece3.9319. PMID: 36248677.

19. Klüttgen B, U Dülmer, M Engels, H.T Ratte. 1994. ADaM, an artificial freshwater for the culture of zooplankton. Water Research 28 (3), 743–746. https://doi.org/10.1016/0043-1354(94)90157-0.

20. Holm, S. 1979. A simple sequentially rejective multiple test procedure. Scandinavian Journal of Statistics. 6 (2): 65–70.

20. Spanier, K.I., Leese, F., Mayer, C., Colbourne, J.K., Gilbert, D., Pfrender, M.E., & Tollrian, R. 2010. Predator-induced defenses in Daphnia pulex: selection and evaluation of internal reference genes for gene expression studies with real-time PCR. BMC Mol Biol, 11, 50. doi: 10.1186/1471-219911-50. PMID: 20587017.

21. Rao X., Huang. X., Zhou, Z., & Lin, X. 2013. An improvement of the 2^(-delta delta CT) method for quantitative real-time polymerase chain reaction data analysis. Biostat Bioinforma Biomath, 3, 3, 71–85. PMID: 25558171.

22. Smirnov, N.N. 2017) Physiology of the Cladocera. 2nd edition Academic Press.

23. Krycer, J.R., Quek, L.E., Francis, D., Fazakerley, D.J., Elkington, S.D., Diaz-Vegas, A., Cooke, K.C., Weiss, F.C., Duan, X., Kurdyukov, S., Zhou, P.X., Tambar, U.K., Hirayama, A., Ikeda, S., Kamei, Y., Soga, T., Cooney, G.J., & James, D.E. 2020. Lactate production is a prioritized feature of adipocyte metabolism. J Biol Chem, 295, 1, 83–98. doi: 10.1074/jbc.RA119.011178. PMCID: PMC6952601.

